# Impacts and interactions of stress, noradrenaline and serotonin signalling on probabilistic reversal learning

**DOI:** 10.64898/2026.07.03.736287

**Authors:** Olivia Stupart, Livia J.F. Wilod Versprille, Katharina Zuhlsdorff, Clara Velazquez-Sanchez, Matthew C.D. Bailey, Judy Chen, Rebecca P. Lawson, Jeffrey W. Dalley

**Author notes:** **Author for correspondence**: Professor Jeffrey W. Dalley, Department of Psychology, University of Cambridge, Downing St, Cambridge CB2 3EB, UK. |. Joint first authors (Olivia Stupart, Livia J.F. Wilod Versprille).

## Abstract

**Rationale:** Early life stress (ELS) is acknowledged to underlie cognitive and emotional abnormalities linked to stress-related mood disorders. ELS can lead to persistent biases in how uncertain feedback is processed to affect the flexibility of decision-making.

**Objectives:** (1) To investigate the effects of ELS on the flexibility of rats trained on a serial probabilistic reversal learning (PRL) task involving spurious positive and negative feedback. (2) To elucidate the involvement of the stress hormone corticosterone and the noradrenergic and serotonergic systems in modulating how ELS affects PRL.

**Methods:** Male and female rats were intermittently separated from maternal care on postnatal days five to nineteen, inclusively. As adults, the same rats were trained on a deterministic reversal learning task involving certain rewarded or non-rewarded outcomes followed by a PRL task where correct and incorrect responses were rewarded on 80% and 20% of trials, respectively. Dose-dependent effects of the beta-blocker, propranolol, selective serotonin reuptake inhibitor, citalopram and corticosterone were subsequently determined.

**Results:** ELS resulted in an increased responsivity to feedback, specifically in males making more win-stay responses following a reward, that was associated with an increased punishment learning rate. In both control and MS rats, propranolol increased feedback sensitivity, but delayed updating following a rule switch. In contrast, neither citalopram nor corticosterone significantly affected reversal learning.

**Conclusions:** ELS is sufficient to cause persistent changes in how feedback is processed by male rats on a reversal learning task. Activation of beta-adrenergic receptors may be necessary for updating learned associations during decision-making involving uncertain feedback.

## Introduction

Early life stress (ELS) is linked to an increased risk for adverse mental health outcomes (Brent and Silverstein 2013; Teicher et al. 2022) with several studies showing ELS to heighten vulnerability to stress-related disorders such as depression and anxiety (Cohen et al. 2006; Nanni et al. 2012; Agnew-Blais and Danese 2016). Impaired uncertainty processing is a proposed mechanism underlying these disorders, leading to maladaptive decision-making (Cella et al. 2010; Wang et al. 2014; Browning et al. 2015). Under stressful conditions, individuals often rely on generalisation to guide decision-making (Schick et al. 2015), but overgeneralisation - particularly to negative feedback – may lead to deficits in uncertainty processing, a feature of depression and anxiety (Elliott et al. 1997, 2002; Lissek 2012).

The widely used procedure of repeated maternal separation (RMS) in rodents attempts to simulate aspects of early life adversity in humans and is recognized as a form of uncontrollable stress that causes persistent behavioural, neural and neuroendocrine abnormalities (Matthews and Robbins 2003; Nishi et al. 2014). Recently, we reported that RMS-induced stress impairs performance on a probabilistic reversal learning (PRL) task, which assesses a subject’s ability optimally to track rewarded outcomes under probabilistic (i.e., uncertain) conditions while disregarding spurious negative feedback (Dutcher et al. 2023). Findings using the PRL task demonstrated that following a second stressor, adults rats deprived of early maternal care were slower to respond to correct target stimuli and showed differential sensitivity to negative feedback compared with control animals (Zühlsdorff et al. 2023; Dutcher et al. 2023). During stress, rodents release corticosterone into the bloodstream, the glucocorticoid equivalent of cortisol in humans (Smith and Vale 2006). Systemic administration of corticosterone in rats has been shown to modulate decision-making under uncertainty by enhancing learning to salient stimuli, reducing negative feedback sensitivity and increasing choice deliberation time after reversals (Bryce and Floresco 2021). However, it remains unclear whether maternal separation stress alone is sufficient to cause persistent biases in the processing of positive and negative feedback.

Noradrenergic and serotonergic neurons originating from the locus coeruleus (LC) and raphe nucleus respectively, are implicated in stress-related disorders related to impaired feedback responsiveness (Soubrié 1986; Itoi and Sugimoto 2010; McCall et al. 2017; Morris et al. 2020; Delcourte et al. 2021; Vahid-Ansari and Albert 2021). Noradrenaline (NA) plays a key role in stress responses and is posited to act as a “reset” signal facilitating new and updated learning in dynamic environments (Yu and Dayan 2005; Bouret and Sara 2005; Avery et al. 2012). NA may thus favour the switch from retrieval of information to encoding (Grella et al. 2021). Accordingly, LC activity is expected to peak during the reversal of a stimulus-reward contingency corresponding with the novelty of new incoming information (Sara et al. 1994). This change in LC activity appears to be adaptive as shown by improved reversal learning following chemogenetic activation of the LC (Rorabaugh et al. 2017). In a complementary manner, serotonin (5-HT) affects how learning proceeds by altering the sensitivity of subjects to positive and negative feedback (Bari et al. 2010). Thus, a low dose of the selective 5-HT reuptake inhibitor (SSRI) citalopram (1 mg/kg) increased lose-shift behaviour on a PRL task (i.e. an increased reaction to negative feedback) whereas a higher dose (10 mg/kg) decreased both lose-shift behaviour and win-stay behaviour, the latter effect an index of reward sensitivity (Bari et al. 2010). However, a follow-up study reported doses of citalopram above 3 mg/kg to increase the proportion of win-stay trials with a non-significant trend decrease in lose-shift behaviour at higher doses (Wilkinson et al. 2020). The basis for this discrepancy in win-stay behaviour to acute higher doses of citalopram is unclear but repeated injections of high-dose citalopram was nevertheless found to increase the frequency of win-stay behaviour on the PRL task (Bari et al. 2010).

The main objectives of the present study were twofold; first, to elucidate whether ELS is sufficient to persistently affect neurobehavioural outcomes measured during adulthood and second, to investigate the relative involvement of the brain’s noradrenergic and serotonergic systems in mediating the behavioural sequelae of ELS.

The neurobehavioural effects of ELS were investigated by comparing the performance of control and maternally separated (MS) rats on both deterministic (i.e., no uncertainty in trial outcomes) and probabilistic variants of a serial reversal learning task. Since MS stress is widely reported to affect the brain’s noradrenergic (Swinny et al. 2010; Hernández-Pérez et al. 2019; Fóscolo et al. 2022) and serotonergic (Llorente et al. 2010; Rentesi et al. 2010; Ohta et al. 2014) systems we hypothesised that MS and control subjects would be differentially sensitive to pharmacological interventions targeting these systems. We first tested the β-adrenergic receptor antagonist propranolol that reduces anxiety (Rex et al. 1998; Zaidi et al. 2020) and increases behavioural flexibility in rats (Hecht et al. 2014). We next tested low and high acute doses of the SSRI citalopram, which we reasoned would differentially affect the sensitivity of animals to negative and positive feedback. Finally, we investigated the effects of the stress hormone corticosterone, an intervention shown previously to affect decision making under uncertainty (Bryce and Floresco 2021).

## Methods

### Subjects

Fourteen timed-pregnant female Lister-Hooded rats were purchased from Envigo (Blackthorn, UK) and delivered on gestational day 14 (E14). After a 1-week habituation period to the animal facility, the pregnant rats littered spontaneously on E21 (designated post-natal day 0 - P0). Within 1 day of birth, pups were sexed using anogenital distance, with males exhibiting a larger anogenital distance than females. Where necessary, cross-fostering was applied to create a minimum of 2 male (M) and 2 female (F) pups per litter. Dams and their litters were then randomly assigned to control or MS groups such that MS males = 16, MS females = 16, control males = 16 and control females = 16. Food was provided *ad libitum* until P56, after which food restriction was imposed to maintain animals at no less than 85% of their age- and sex-adjusted predicted weights. Water was provided *ad libitum* throughout the study. The temperature of home cages was maintained between 20-22°C with a relative humidity of 55 ± 3%. Dams and litters were kept on a 12h/12h light/dark cycle with lights on at 07:00. Post-weaning pups were subsequently housed and remained for the duration of the experiment, under a reverse light cycle with lights off at 07:00. Experiments were conducted in accordance with Project Licence PA9FBFA9F and the UK Animals (Scientific Procedures) Act 1986 (amendment regulations 2012) as well as the EU legislation on the protection of animals used for scientific purposes (Directive, 2010/63/EU) following ethical review by the University of Cambridge Animal Welfare and Ethical Review Body (AWERB).

### Maternal separation procedure

Aside from routine husbandry, control litters were left undisturbed until weaning (P21). They were only exposed to standard animal facility care with minimal handling aside from twice-a-week cage cleaning and ear notching for identification on P12. MS litters were separated from their dam from 09:00 to 15:00 during the ‘lights-on’ period. Maternal separation was repeated daily from P5 to P19, inclusively (Fig.1A). Each litter was moved, in the same order each day, by counting out the pups into a smaller litter-specific cage with littermates kept together. The separation cage was 24 x 40 x 19 cm in size and had paper bedding and sawdust with a red Perspex lid. Food and water were available *ad libitum*. Separation cages were moved to a different room and placed inside a ventilated surgical incubator kept at 30-32°C. Pups were monitored for signs of dehydration hourly. No adverse health impacts were noted.

**Fig. 1.**
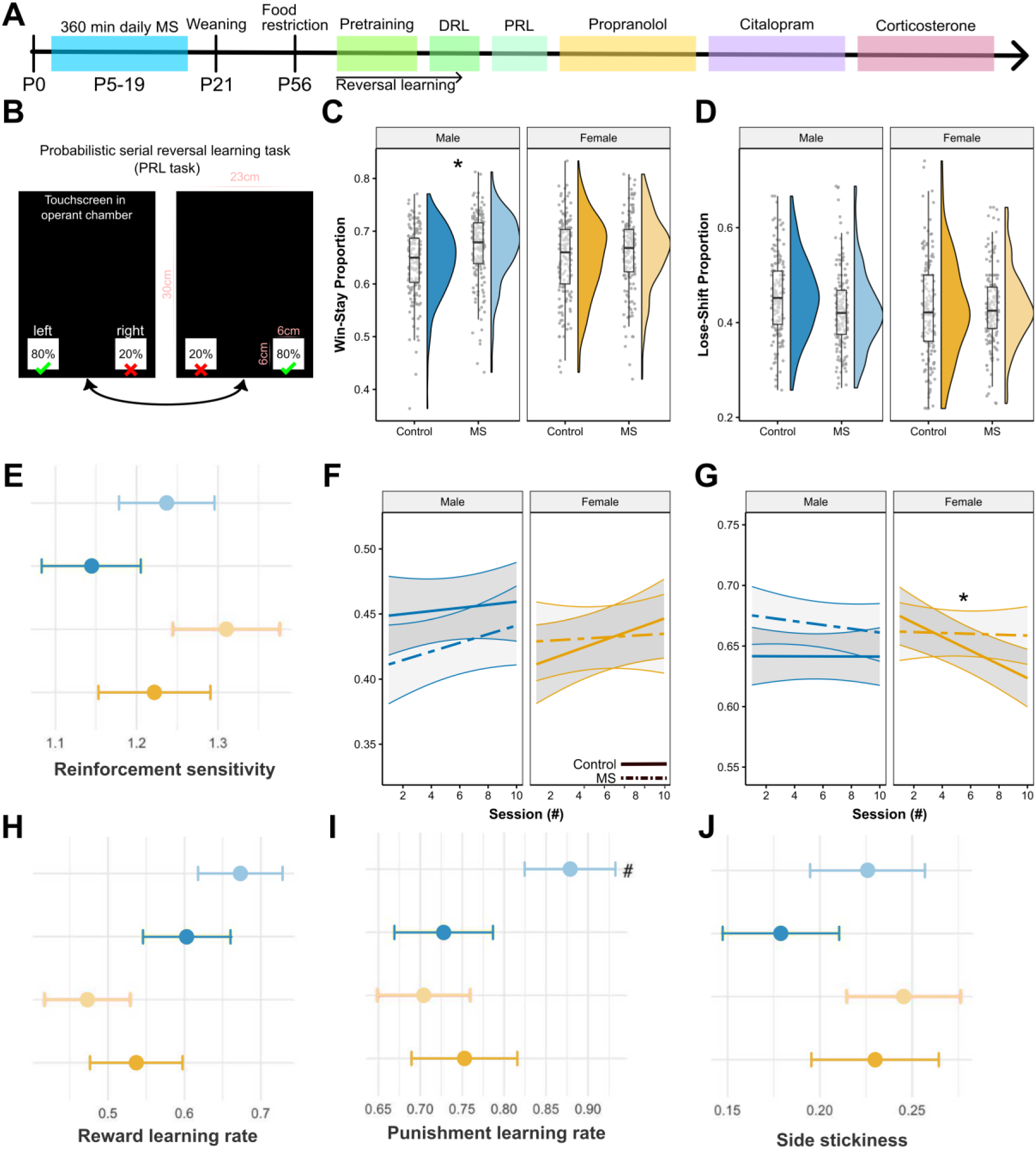
Timeline of experimental procedures (A). Overview of the PRL task (B). Contingencies switch after 8 consecutive correct responses, with 80% reward probability for a correct choice and 20% reward probability for an incorrect choice. Effects of sex (colour) and group (solid vs dashes lines) on PRL performance. Violin and box plot for win-stay probability (C) and lose-shift probability (D). Emtrends plot with 95% CIs for change in win-stay probability (F) and lose-shift probability (G), over consecutive sessions. Results from the hierarchical Bayesian winning RL model, showing differences in group and sex mean parameters on the PRL (E, H, I, J). * Represents p<0.05 significant change over consecutive sessions within the indicated group, # indicates 0 ∉75% HDI.

Dams were kept in the home-cage for the duration of the separation procedure. Reintroduction was conducted cage by cage, in the same order as separation. Pups were massaged in the dams bedding to transfer her scent to their fur, counted and placed back into the home cage. Dam-pup interactions were monitored for several minutes to identify possible aggression or indications of pup rejection. No adverse reintroductions occurred, and nursing was resumed within 5 min for all litters following separation.

### Behavioural testing

Behavioural testing was conducted in two, sex-segregated rooms; each containing eight identical operant chambers (Med Associates, St. Albans, VT, USA) enclosed in a fan-ventilated, sound, and light attenuating box. K-Limbic software (Conclusive Marketing LTD., Herts, UK) was used to control the chambers and touchscreens.

The training protocol is described by (Dutcher et al. 2023). Rats were trained or tested once a day for 5-6 days a week, conducted during the dark period. Sessions ended after 40 min or the completion of 200 trials, whichever came first. After habituation to the operant chambers, training on the PRL task commenced (Fig.1A-B). In the first stage, a 2 x 2 cm visual white square stimulus was randomly presented on the left or right corner of the touchscreen. A response at the stimulus with a nose poke was rewarded with a 45 mg sucrose pellet (TestDiet, St. Louis, MO, USA) indicated by the illumination of the magazine light. The inter-trial interval (ITI) was set to 5 sec and rats were required to make at least 80 responses during every session. The task was identical for the next stage except for the loss of reward for screen touches outside of the stimulus. A punishment in the form of a time out (TO) of 5 sec followed background touches and was reinforced by the illumination of the house light. Subjects were required to make a head entry into the empty food magazine to self-initiate the next trial. It was also a prerequisite for animals to achieve at least 100 correct touches over two consecutive sessions to progress to the next stage. The next stage involved deterministic reversal learning (DRL). Two visually identical stimuli (2 x 2 cm white squares) were simultaneously presented in the left and right bottom corners of the touchscreen. “Correct” and “incorrect” stimuli were randomly allocated for each subject. Responses on the “correct stimulus” were rewarded as before. Responses to the “incorrect” stimulus as well as background touches were punished in the same way as omissions (i.e., a failure to respond within 5 sec). Eight correct responses in a row resulted in the reversal of the stimulus-reward contingency. Subjects were required to achieve 4 reversals during two consecutive sessions to proceed to the final PRL stage. The PRL task differed from the DRL, where rewards were only delivered following 80% of responses to the “correct” side, and following 20% of responses to the “incorrect” side. Animals were thus required to disregard the spurious feedback that occurred on 20% of trials. A reversal of the stimulus reward contingency occurred following 8 consecutive correct responses.

### Pharmacological interventions

The effects of pharmacological agents on the behavioural outcomes of the PRL were investigated using counterbalanced, within-subject, Latin square designs. Each drug administration session was preceded by a baseline session and followed the next day with no training. A washout period of at least 3 days was imposed between different drugs. Treatment groups were allocated in a pseudo-randomised design to ensure equal distribution of the sex and MS conditions. All subjects received every dose of each drug, except for four females, who had delayed task acquisition and therefore did not receive propranolol, but were included in the citalopram and corticosterone studies. All pharmacological agents were systemically administered with a volume of 1 ml/kg *via* the intraperitoneal route (Terumo® Agani™ 0.5×16 mm needles and Terumo® 1 ml syringes) 30 min prior to testing. Pharmacological agents were dissolved in 0.9% sterile saline (Animalcare®, UK) unless otherwise specified.

The β-blocker propranolol (Sigma-Aldrich, UK, P0884) (Hauser et al. 2017) was administered in the following doses: 1, 3 and 10 mg/kg, in accordance with prior studies (Chiamulera et al. 2010; Rodberg et al. 2023). The SSRI citalopram (Sigma-Aldrich, UK, C7861) (Wong et al. 1991; Richelson 1994) was given in the following doses: 1, 3 and 10 mg/kg, in accordance with prior studies (Bari et al. 2010; Wilkinson et al. 2020). The glucocorticoid, corticosterone (Cayman Chemicals, Ann Arbor MA, USA, 50-22-6), was given in the following doses: 0.5, 5 and 10 mg/kg, in accordance with previous studies (Munera et al. 2017; Ossenkopp et al. 2011). Corticosterone was first dissolved in 40% HP-β-cyclodextrin (Boehringer Ingelheim, Germany), sonicated in a water bath at 35°C for 2 hours, adjusted to a pH of 7.4-7.6, then diluted to 20% in 0.9% saline. Each Latin square design included the appropriate vehicle; 0.9% saline for propranolol and citalopram, and 20% HP-β-cyclodextrin in 0.9% saline for corticosterone.

### Data and statistical analysis

Data was exported for preprocessing with custom scripts in Jupyter (version 7.0.8) with subsequent data presentation and statistical analysis carried out using RStudio (version 4.3.2). Data were initially assessed for normality and transformed where appropriate. All analyses used mixed-effect linear models to allow for repeated measures within subjects, with the following fixed effects: sex, MS or control group and drug dose. Random effects were a combination of subject, operant chamber ID or session.

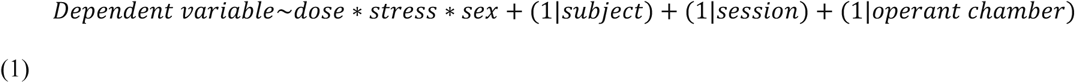

Random effect structures were selected for each behavioural outcome using the first 10 sessions of PRL testing, selecting the most appropriate model using the likelihood ratio test and using AIC and BIC where the optimal model was unclear. This resulted in independent model structures for each behavioural outcome, but the same model was used when repeating the analysis for each training stage or drug challenge, with these random effects structures selected: none for the number of perseverative responses (square root transformation); subject, session and operant chamber for response latency (logarithmic transformation); subject and operant chamber for collection latency (logarithmic transformation); subject and session for accuracy, number of trials and initiation latency (logarithmic transformation); subject for reversal rate, number of trials to first reversal (square root transformation), win-stay probability, lose-shift probability, and ‘true’ win-stay probability.

Mixed linear model assumptions were inspected visually using the *performance check_model* package and function in R. *Post-hoc* pairwise comparisons using emmeans and emtrends (Lenth 2024) were completed only where a main effect or interaction was identified, with appropriate Bonferroni adjustments for multiple comparisons. Comparisons used estimated marginal means (EMMs), which are model-adjusted mean values for each group. Significant pairwise differences are reported with t-statistics and p-values. Emtrends predicted the model-adjusted gradient of effect for successive sessions of testing on a continuous scale. Emtrends are reported when the gradient was significantly different from 0 (trend and p-value) and when the gradients were significantly different between groups (t-statistic and p-value). Test statistics and *p*-values are provided in the main text for all significant (*p<0.05*) effects involving group or drug, significant effects of sex are reported in the supplementary online materials section. Means are reported with SEM following in brackets. Figures display median and spread of the raw data, with factor adjusted EMMs labelled where appropriate, even if transformed data were used for the models.

### Hierarchical Bayesian reinforcement learning modelling

Reinforcement learning models were fit to the reversal learning data using a hierarchical Bayesian approach. Three different models with different parameters were implemented in Stan (version 2.26.1). The highest hierarchical level incorporated a group-specific mean and standard deviation for each parameter. A summary of the priors set for each parameter are given in Table 1. A normal distribution with the respective mean and standard deviation was used to select the parameters for each subject. Subsequently, a reinforcement learning model was fitted to the behaviour, and the highest posterior density interval (HDI) was calculated for group mean differences.

**Table 1.** Priors for model parameters.

| Parameter | Prior | Reference |
| --- | --- | --- |
| Reward learning rate, $\alpha_{rew}$ | Beta(1.2, 1.2) | (den Ouden et al. 2013) |
| Punishment learning rate, $\alpha_{pun}$ | Beta(1.2, 1.2) | (den Ouden et al. 2013) |
| Combined learning rate, $\alpha$ | Beta(1.2, 1.2) | (den Ouden et al. 2013) |
| Reinforcement sensitivity, $\beta$ | Gamma($\alpha=4.82$,<br>$\beta=0.88$) | (Gershman 2016) |
| Side stickiness, $\kappa_{side}$ | Normal(0,1) | (Christakou et al. 2013) |
| **Intersubject variability in parameters** |  |  |
| $\alpha_{rew}$, $\alpha_{pun}$, $\alpha$, $\kappa_{side}$ intersubject standard deviations | Normal(0,0.05)<br>constrained to $\geq 0$ | (Kanen et al. 2019) |
| $\beta$ intersubject standard deviations | Normal(0,1) | (Gershman 2016) |

Q-values were updated on each trial using the following equation:

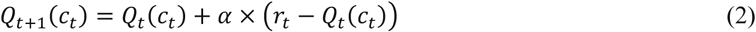

where *Qt*+1(*ct*) is the Q-value of the stimulus selected on the current trial for the next trial, *Qt*(*ct*) is the expected value of the chosen stimulus on the current trial, *α* is the learning rate (a higher rate being reflective of faster learning) and *rt* is the reinforcement on trial *t* (1 for reward and 0 for punishment). In the next step, the softmax decision rule was used to calculate the probability of making one of two choices:

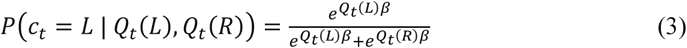

Qt(L) and Qt(R) are the Q-values of the left and right stimuli, and β is the reinforcement sensitivity parameter, also known as the exploitation vs exploration parameter. Lower values of β indicate greater exploration and lower sensitivity to reinforcement, whereas greater values represent increased exploitation and greater sensitivity to reinforcement.

The models that were fitted to the data were the following:

1. **Two parameters: α and β**, the learning rate and reinforcement sensitivity parameter.
2. **Three parameters: α, β,** side stickiness parameter **κside**, which represents the tendency to choose the same side as on the previous trial. It was used to update the Q-values using the following equation: *Q^loc^_s_*_,*t*_ = *κ_side_L_s_*_,*t*−1_. *L_s_*_,*t*−1_ is the side chosen by the subject on the previous trial. If the same side was chosen this value is equal to 1, and 0 if the other side was chosen. The final Q-value is the sum of *Q^loc^_s_*_,*t*_ and the Q-value as calculated in equation 1.
3. **Four parameters: αrew, αpun, β and κside.** In this model, the combined learning rate was separated into two learning rates for reward and punishment.

Hamiltonian Markov chain Monte Carlo sampling was used to fit the models via Stan 2.17.2 (Carpenter et al. 2017). The potential scale reduction factor was used to ensure convergence. A value close to 1 indicates perfect convergence. A cut-off of 1.1 was selected as a stringent criterion for convergence. Models were compared using a bridge sampling estimate of the marginal likelihood using the “bridgesampling” R package (Gronau et al. 2017).

## Results

To investigate the effects of ELS on reversal learning and feedback, the performance of MS and control rats was assessed on the DRL and PRL tasks. This analysis revealed that MS did not significantly affect responding to deterministic feedback following a reversal in the stimulus-reward contingency. When a probabilistic reward contingency was introduced, all subjects, irrespective of MS or sex, changed their responses to feedback (see Table 2). Thus, introducing probabilistic feedback resulted in reduced accuracy across all subjects and an increased latency to respond.

**Table 2.** Summary of mean (SD) values for behavioural outcomes during the last 5 DRL training sessions, first 10 PRL sessions, and overall combined values. Control and MS groups are collapsed (n = 64).

| Behavioural Measure | Combined |  | DRL |  | PRL |  |
| --- | --- | --- | --- | --- | --- | --- |
|  | Mean | SD | Mean | SD | Mean | SD |
| % Correct | 0.65 | 0.09 | 0.66 | 0.1 | 0.65 | 0.07 |
| No. Trials | 130.1 | 32.7 | 124.0 | 36.9 | 135.6 | 27.3 |
| No. Reversals | 4.71 | 2.89 | 5.29 | 3.37 | 4.18 | 2.27 |
| Reversal Rate | 0.03 | 0.02 | 0.04 | 0.02 | 0.03 | 0.01 |
| No. to 1<sup>st</sup> Rev. | 28.6 | 19.8 | 24.2 | 14.0 | 32.8 | 23.4 |
| Mean Perseverations | 1.93 | 1.12 | 2.06 | 1.29 | 1.80 | 0.91 |
| Mean Response Latency | 364.4 | 117.6 | 391.9 | 131.3 | 339.2 | 97.8 |
| Mean Collection Latency | 139.3 | 64.4 | 162.5 | 71.1 | 118.1 | 49.1 |
| Mean Initiation Latency | 702.1 | 812.4 | 834.2 | 1038.4 | 581.5 | 506.4 |
| Win-Stay Proportion | 0.71 | 0.09 | 0.74 | 0.08 | 0.69 | 0.09 |
| Lose-Shift Proportion | 0.43 | 0.11 | 0.45 | 0.13 | 0.40 | 0.09 |

With respect to PRL performance measures, the number of trials completed, percentage accuracy, reversal rate and the number of trials to the first reversal were unaffected by MS. There was also no main effect of MS on the response latencies to choose, collect reward or initiate the next trial. However, a significant interaction between stress × sex was revealed for win-stay responding (F_(1,219)_ = 4.06, p = 0.045; Fig.1C) and “stay” responses following a true win (StTW) (F_(1,233)_ = 3.96, p = 0.048), which did not extend to lose-shift responding (Fig.1D). Pairwise comparisons of estimated marginal means (EMM) between MS and controls for each sex indicated there was no significant difference between groups in females for win-stay or StTW. Males MS subjects, however, made significantly more “stay” responses after a win (t = 2.03, p = 0.047), specifically after a true win (t = 1.96, p = 0.05) compared with control males.

After comparison of the three RL models described in the methods section, it was found that Model 3 was the best fitting model (Table 3). This model included the parameters **α_rew_** (reward learning rate, **α_pun_** (punishment learning rate), **β** (reinforcement sensitivity) and **κ_side_** (side stickiness parameter). Analysis with this model also demonstrated sex-stress-specific differences in feedback sensitivity. Thus, there was an increase in punishment learning rate **α_pun_** for MS males compared to control males (Fig.1I), group difference 75% highest density interval (HDI), but no difference in reward learning rate **α_rew_** (Fig.1H), reinforcement sensitivity **β** (Fig.1E) or **κ_side_,** side stickiness (Fig.1J). There were no significant interactions for any of the other behavioural variables. Therefore, MS did not affect how subjects updated their responding to the PRL task. However, when including sex, there was a significant interaction between sex x session number x stress group for the win-stay responding (F_(1,572)_ = 4.04, p = 0.045) and StTW (F_(1,572)_ = 4.66, p = 0.031). This was driven by a specific reduction in win-stay responding and StTW across sessions for female control subjects (Fig.1F-G).

**Table 3.** Model comparison summary. Model 3 achieved the highest marginal likelihood in all treatment groups and overwhelmingly dominated the model posterior distribution, indicating that asymmetric learning from positive and negative outcomes provides the best account of behaviour.

| Model | Parameters | Rank (all) | Log marginal likelihood (propranolol) | Log posterior P ( propranolol ) | Log marginal likelihood (citalopram) | Log posterior P (citalopram) | Log marginal likelihood (corticosterone) | Log posterior P (corticosterone) |
| --- | --- | --- | --- | --- | --- | --- | --- | --- |
| 1 | $\alpha, \beta$ | 3 | -20933 | -3381.41 | -19336 | -153.20 | -17494 | -93.92 |
| 2 | $\alpha, \beta, \kappa_{stim}$ | 2 | -20754 | -3344.88 | -19288 | -104.88 | -17450 | -49.98 |
| 3 | $\alpha_{rew}, \alpha_{pun}, \beta, \kappa_{side}$ | 1 | -20629 | 0.000 | -19183 | -0.000 | -17400 | 0.000 |

### Propranolol increases repetitive responding on a probabilistic reversal learning task

Significant main effects on reversal rate (F_(4,217.8)_ = 6.17, p = 0.0001; Fig.2A) and mean perseverative responses (F_(4,262)_ = 2.55, p = 0.04; Fig.2B) were revealed following the administration of propranolol. The rate of reversals is generally regarded as a robust measure of task performance, with an increased rate indicative of fast, adaptive learning of current stimulus-reward contingencies. Under propranolol, the reversal rate increased up to a maximum at 3 mg/kg, which was significantly greater than vehicle (t = 4.39, p = 0.0002), and the baseline mean (t = 3.86, p = 0.0015). The number of perseverative responses was inversely related to the speed at which subjects update rule learning following a stimulus-reward contingency reversal. The mean number of perseverations was significantly increased in response to 10 mg/kg propranolol when compared to 1 mg/kg (t = 2.89, p = 0.042). Subjects therefore achieved more reversals per session overall, but immediately following a reversal perseverated in their responding for longer. These effects are consistent with a global increase in repetitive responding such that when a new rule is learnt there is less variability in responding despite the new rule being learnt more slowly. In relation to feedback sensitivity on the PRL task, win-stay responding increased following a dose of 3 mg/kg compared with vehicle (t = 4.14, p=0.0005; Fig.2C) and the 10 mg/kg dose level (t = −3.46, p = 0.0065), which is consistent with a global increase in stickiness. Furthermore, lose-shift responding decreased compared with vehicle and the 3 mg/kg dose level (t = −3.12, p = 0.0208; Fig.2D). Thus, the repetitive responding caused by propranolol was independent of feedback valence and reflected instead a general reduction in choice variability.

**Fig. 2.**
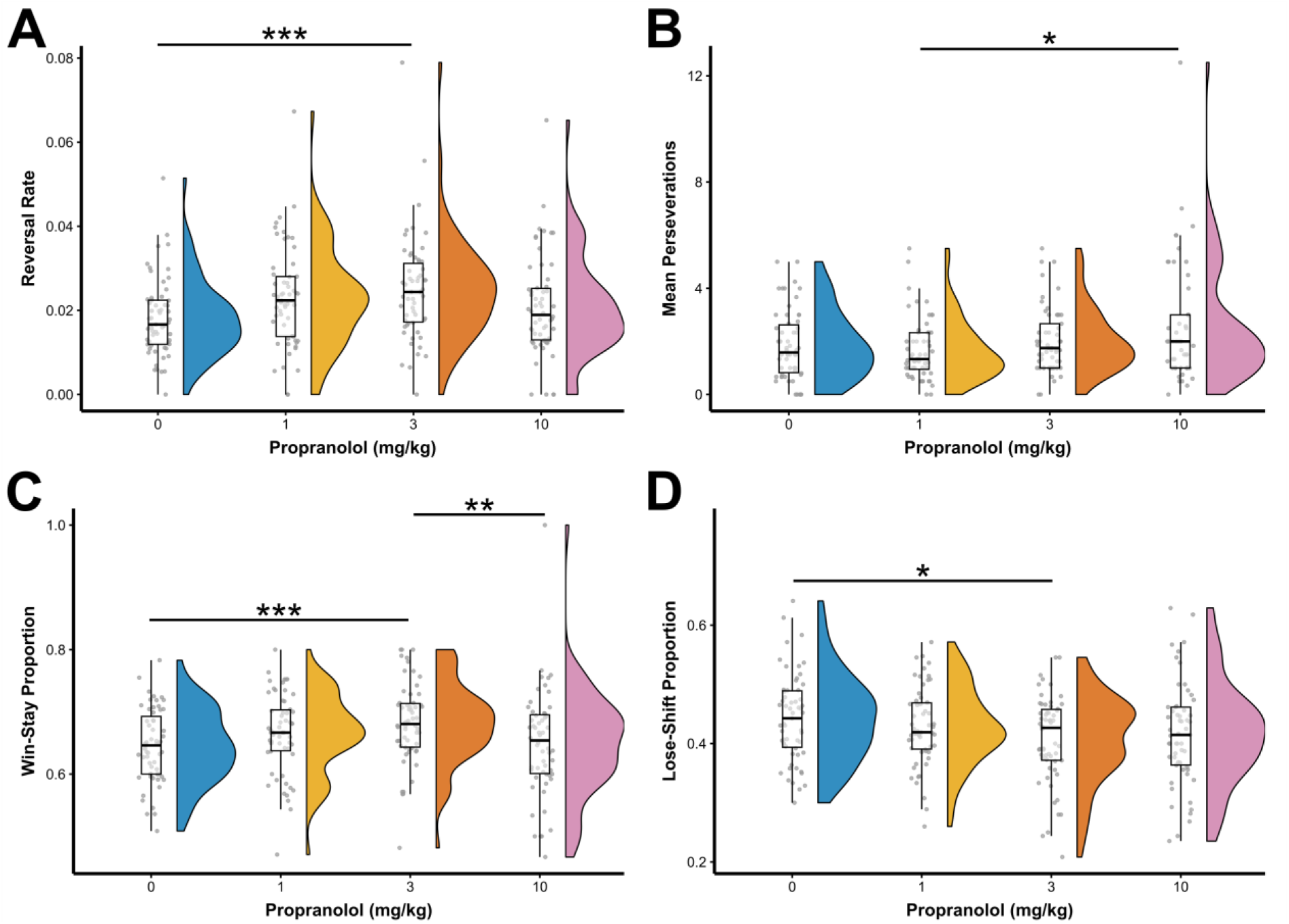
Violin and box plot for reversal rate (A), perseverative responses (B), win-stay proportion (C) and Lose-Shift proportion (D) of all subjects following the administration of propranolol. There was a dose dependent increase in reversal rate with the administration of propranolol to a maximum of 3mg/kg. This was driven by both by an increase in win-stay responding, and a decrease in lose shift responding, such that at 3mg/kg subjects are least likely to change their response, regardless of feedback. There was a dose dependent increase in the mean perseverations after a change in choice contingency.

The administration of propranolol was accompanied by a significant reduction in the number of trials completed in each session (F_(4,32.6)_ = 17.70, p < 0.0001) and increased latencies to respond (F_(4,29.4)_ = 18.13, p < 0.0001), collect food reward (F_(4,218.7)_ = 6.59, p < 0.0001) and initiate the next trial (F_(4,36.7)_ = 4.34, p = 0.006). However, propranolol had no significant effect on the accuracy of responding (F_(4,13.1)_ = 2.54, p =0.09) and significant effects on response latencies were only observed at the highest dose tested (10 mg/kg). Since the effects of propranolol on reversal rate, win-stay, and lose-shift were greatest at 3 mg/kg it is thus unlikely that overt sedation contributed to the effects of propranolol on PRL.

### Lack of effects of citalopram and corticosterone on probabilistic reversal learning

Neither citalopram (Fig.3) nor corticosterone (Fig.4) significantly affected the rate of reversals, number of perseverative responses, or the feedback sensitivity measures of win-stay and lose-shift probability. However, the administration of citalopram was associated with a reduction in the number of trials completed (F_(4,46.9)_ = 11.06, p < 0.0001) and a reduction in the latencies to respond and initiate the next trial (Respond: F_(4,34.4)_ = 3.07, p = 0.029; Initiate: F_(4,48.9)_ = 9.26, p < 0.0001). Corticosterone significantly decreased the latency to collect food reward (F_(4,240.5)_ = 4.15, p = 0.003). This main effect interacted with early life stress (F_(4,240.5)_ = 4.34, p = 0.002) and was only significant for comparisons within the MS group (males and females combined): vehicle versus 0.5 mg/kg (t = −4.69, p < 0.0001); vehicle versus 5 mg/kg (t = −4.37, p = 0.0002). MS animals were thus faster to collect food reward following corticosterone administration, an effect that was not present in control animals administered corticosterone.

**Fig. 3.**
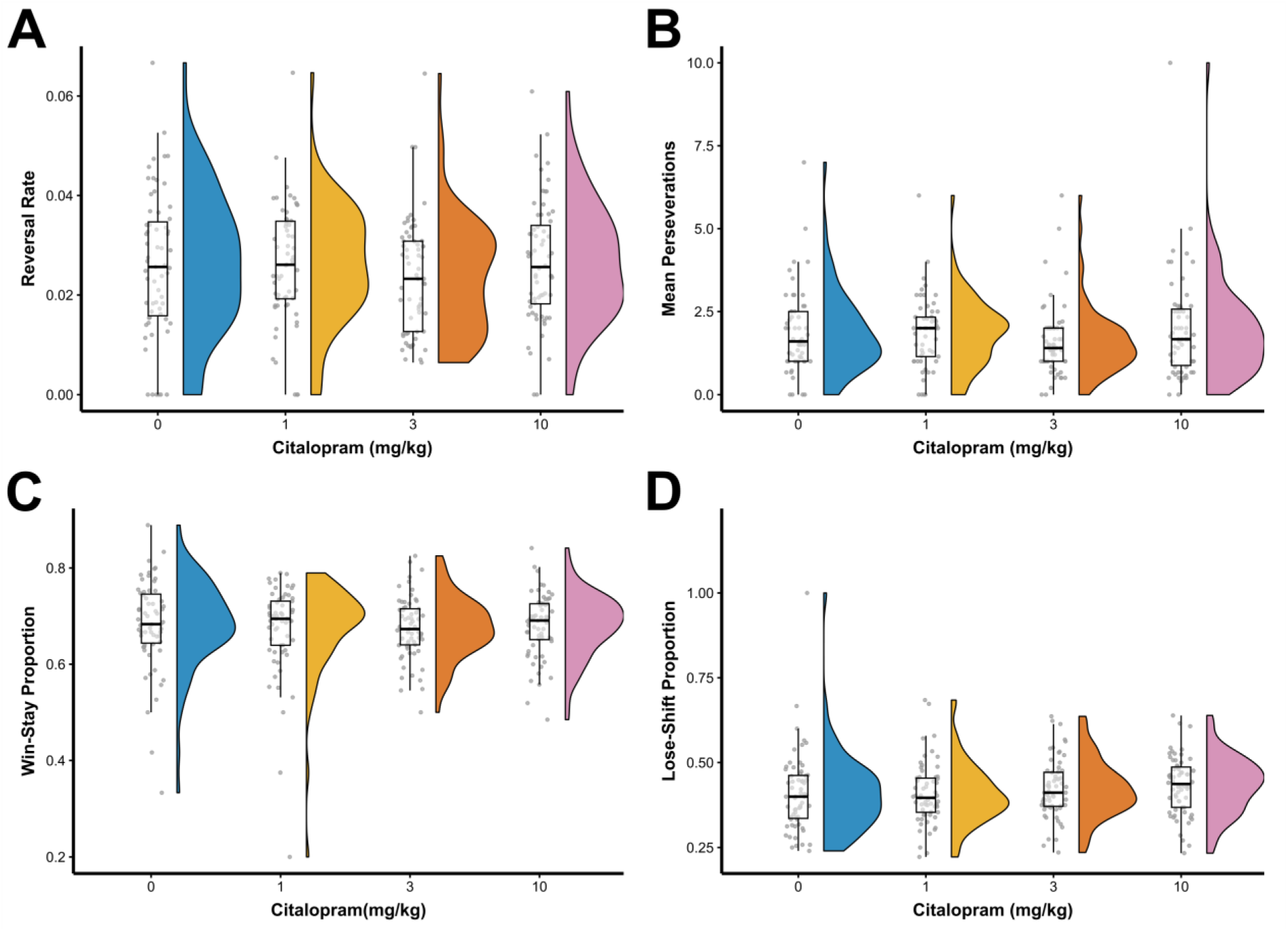
Violin and box plot for reversal rate (A), perseverative responses (B), win-stay proportion (C) and Lose-Shift proportion (D) of all subjects following the administration of citalopram. Citalopram did not produce significant effects across any of the behavioural measures in this task.

**Fig. 4.**
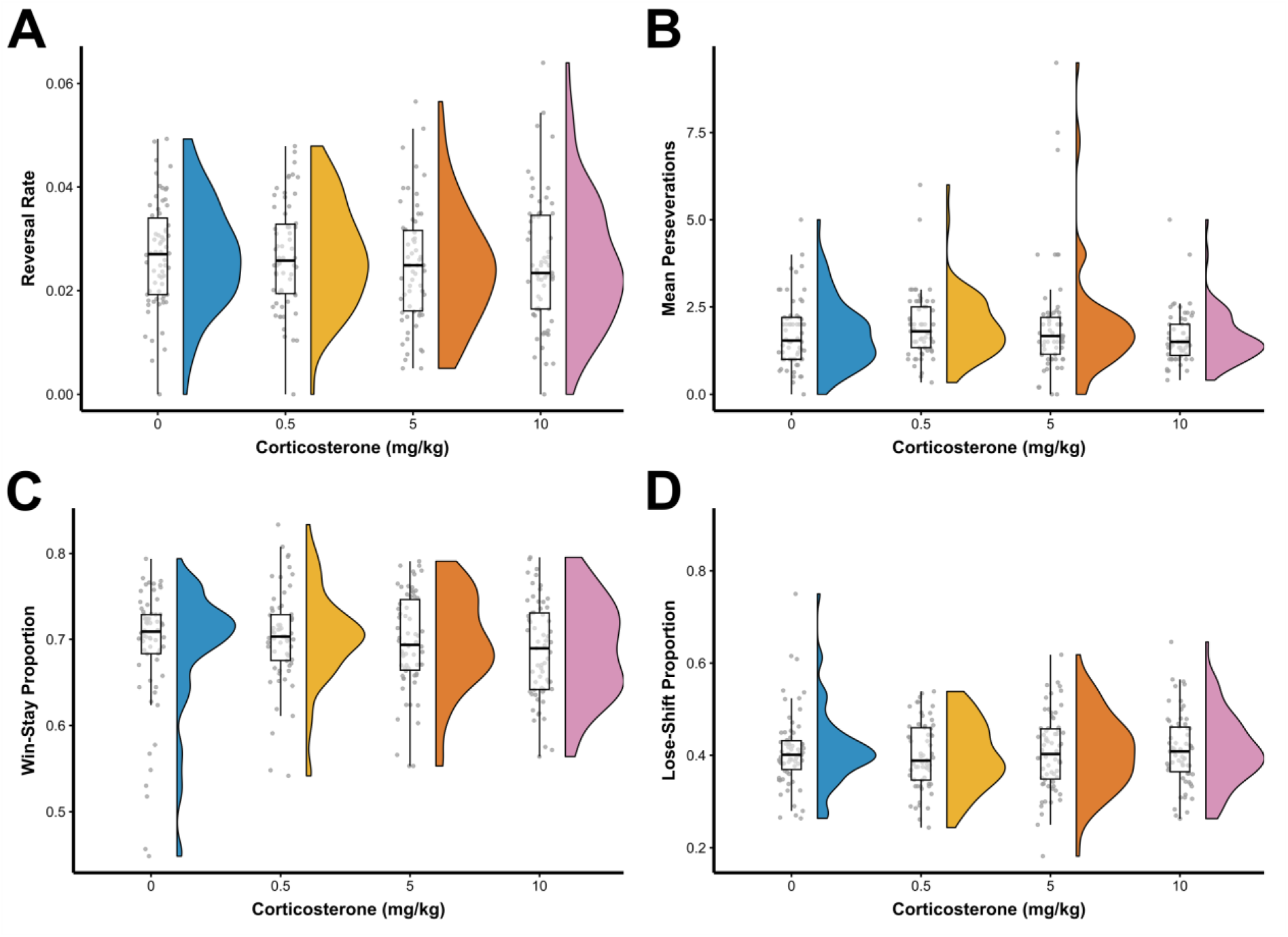
Violin and box plot for reversal rate (A), perseverative responses (B), win-stay proportion (C) and Lose-Shift proportion (D) of all subjects following the administration of corticosterone. Corticosterone did not produce significant effects across any of the behavioural measures in this task.

### Computational modelling

Further comparison of groups using the winning model (model 3) showed that the reward learning rate **α_rew_** significantly decreased following 3 and 10 mg/kg propranolol in both control and MS groups (difference in parameter per-group mean, posterior 95% HDI excluding zero (group difference, 0 ∉ 95% HDI) (Fig.5A-B). High dose (10 mg/kg) propranolol also decreased the punishment learning rate (Fig.5C-D) in the control (group difference, 0 ∉ 95% HDI) and MS groups (group difference, 0 ∉ 75% HDI). Moreover, the reinforcement sensitivity parameter increased in both control and MS groups following 3 and 10 mg/kg propranolol compared to vehicle (group difference, 0 ∉ 95% HDI; Fig.5E-F). However, more restricted effects were evident for the side stickiness parameter with an increased stickiness in the control group across all doses (group difference, 0 ∉ 95% HDI), with no significant effects of propranolol in the MS group (Fig.5G-H).

**Fig. 5.**
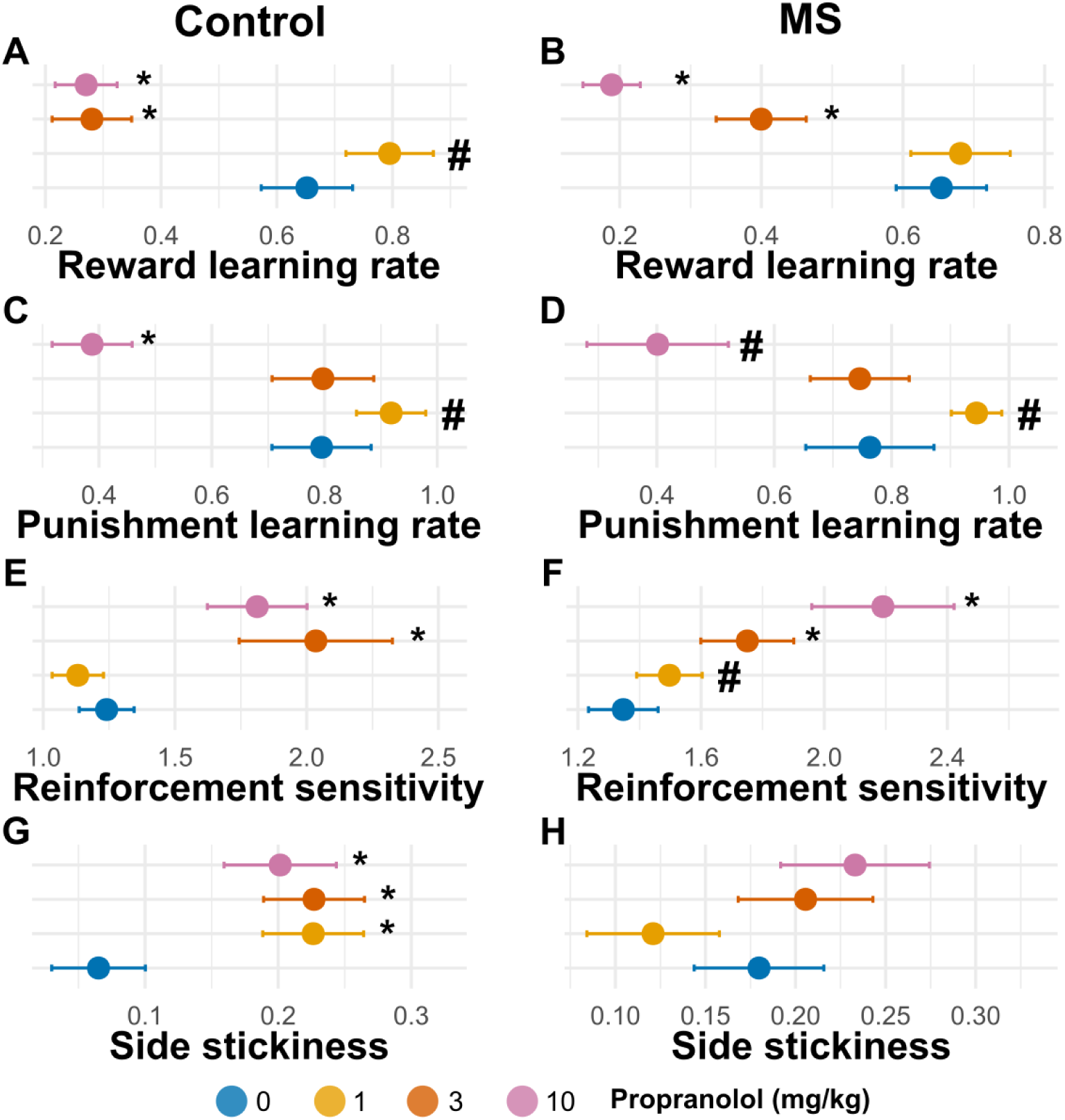
Results from the hierarchical Bayesian winning RL model, showing differences in group mean parameters following the administration of propranolol. (MS, maternal separation; HDI, highest posterior density interval. *Indicates 0 ∉ 95% HDI; # indicates 0 ∉75% HDI). In control animals, propranolol reduced reward and punishment learning rates, increased reinforcement sensitivity, and increased side stickiness relative to vehicle. In MS animals, propranolol reduced reward and punishment learning rates and increased reinforcement sensitivity, while having comparatively little effect on side stickiness.

The same model, Model 3 (**α_rew_, α_pun_**, **β**, **κ_side_**), was the winning model for the citalopram study (Fig.6). In contrast to propranolol, citalopram had no effect on the reward learning rate (Fig.6A-B) and side stickiness (Fig.6G-H). At a dose of 1 mg/kg, citalopram increased the punishment learning rate in both control and MS groups (group difference, 0 ∉ 75% HDI; Fig.6C-D) whereas reinforcement sensitivity decreased in both groups for all doses tested (group difference, 0 ∉ 75% HDI; Fig.6E-F). Model 3 was also the winning model for the corticosterone study (Fig.7). The reward learning rate (Fig.7A-B) was increased in the control group for 10 mg/kg corticosterone (group difference, 0 ∉ 95% HDI) but decreased at all doses in the MS group (group difference, 0 ∉ 95% HDI for both 0.5 and 5 mg/kg, group difference, 0 ∉ 75% HDI, for 10 mg/kg corticosterone). At a dose of 0.5 mg/kg, corticosterone decreased the punishment learning rate in the control (group difference, 0 ∉ 95% HDI), but not in the MS group (Fig7C-D). Reinforcement sensitivity (Fig.7E-F) increased at all doses in the MS group (group difference, 0 ∉ 95% HDI) but decreased at the highest dose (10 mg/kg) in the control group (group difference, 0 ∉ 95% HDI). Side stickiness (FigG-H) increased after the 0.5 mg/kg dose in the control group (group difference, 0 ∉ 75% HDI) but decreased after 10 mg/kg in the MS group (group difference, 0 ∉ 95% HDI).

**Fig. 6.**
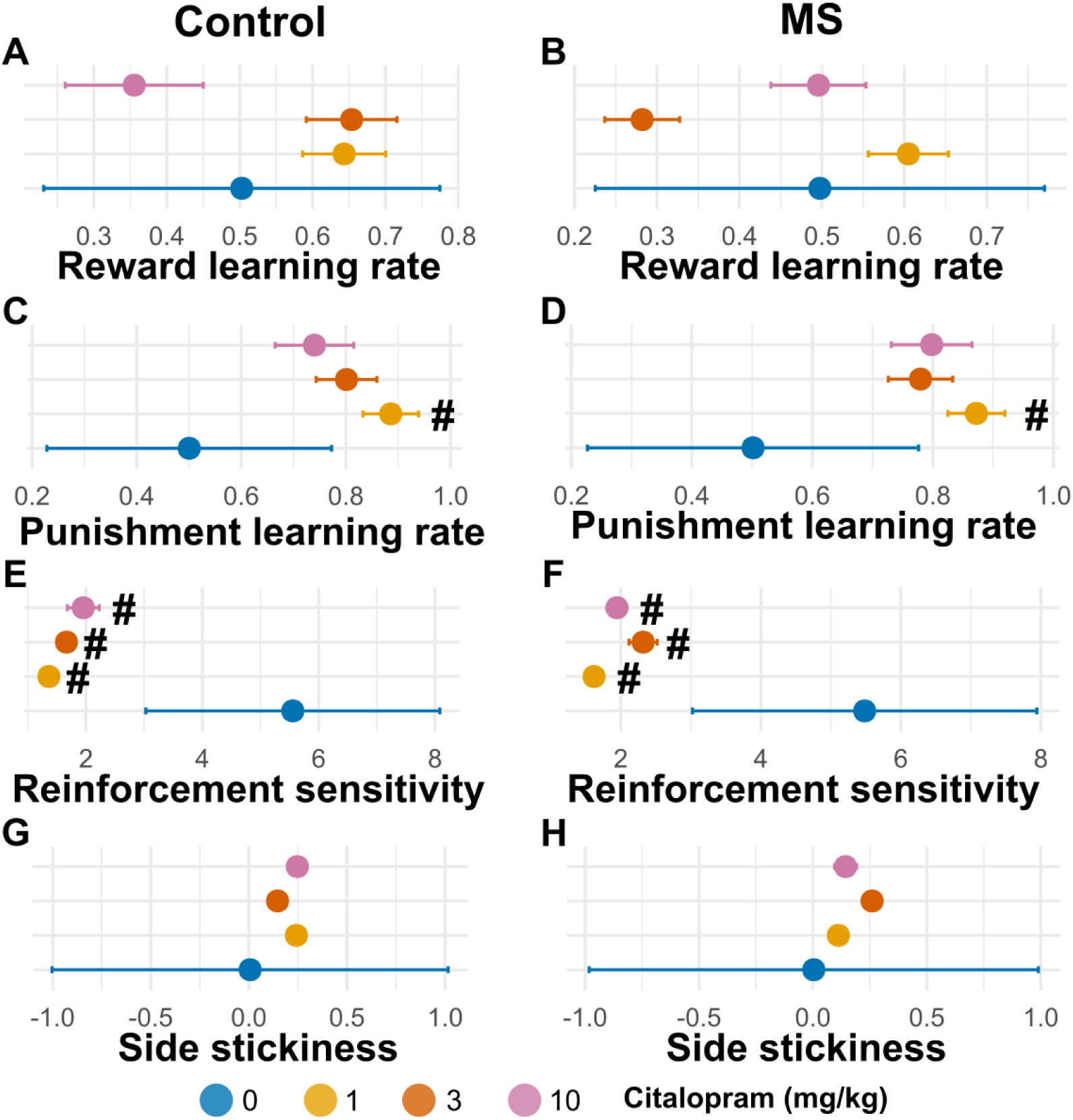
Results from the hierarchical Bayesian winning RL model, showing differences in group mean parameters following the administration of citalopram. (MS, maternal separation; HDI, highest posterior density interval. *Indicates 0 ∉ 95% HDI; # indicates 0 ∉75% HDI). In control and MS animals, low-dose citalopram enhanced punishment learning rate and decreased reinforcement sensitivity across all doses relative to vehicle. Neither reward learning rate nor side stickiness was affected by citalopram.

**Fig. 7.**
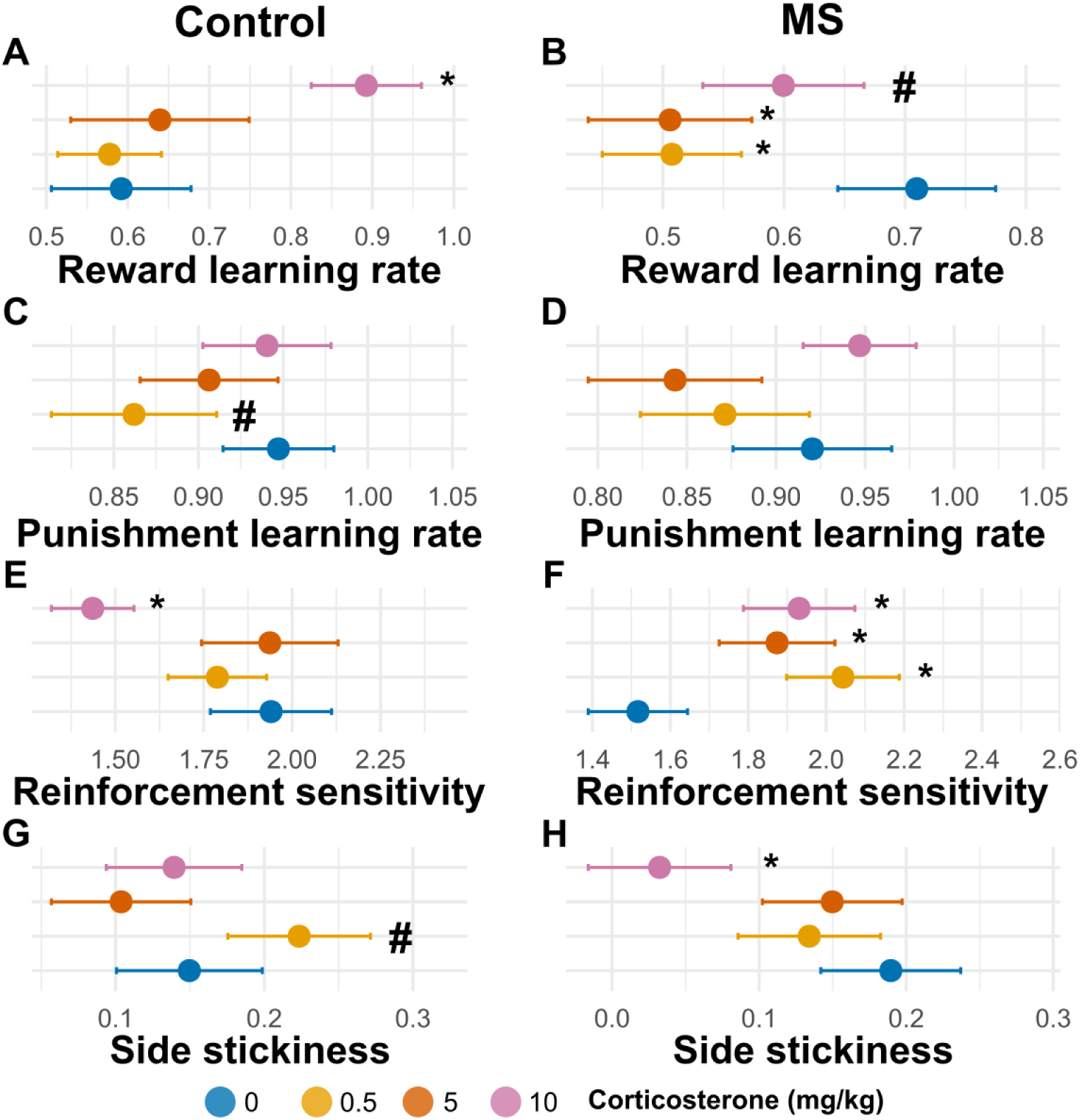
Results from the hierarchical Bayesian winning RL model, showing differences in group mean parameters following the administration of corticosterone. (MS, maternal separation; HDI, highest posterior density interval. *Indicates 0 ∉ 95% HDI; # indicates 0 ∉75% HDI). In control animals, high-dose corticosterone increased reward learning rate while reducing reinforcement sensitivity, while low-dose reduced punishment learning rate and side stickiness. In MS animals, corticosterone reduced reward learning rate and enhanced reinforcement sensitivity. Side stickiness was selectively reduced following high-dose corticosterone. Punishment learning rate remained unaffected in MS animals.

## Discussion

We investigated how ELS in the form of repeated intermittent maternal separation affects how the noradrenergic and serotonergic systems modulate serial reversal learning. Two variants of the reversal learning paradigm were employed: a deterministic task involving a fixed reward probability of 100% on correct trials, and a probabilistic task where reward was delivered on 80% of correct trials and 20% of incorrect trials. Our findings indicate that while MS does not affect reversal learning when reward probabilities are fixed it does affect feedback sensitivity on a PRL task in a sex dependent manner. Specifically, we found that win-stay probability and punishment learning increased in male MS rats compared with male control rats. This effect was relatively selective with no ancillary effects on other behavioural measures, including the rate of reversal. Our pharmacological findings revealed that propranolol robustly affects reversal rate, perseveration and feedback-related responding, whereas citalopram and corticosterone had limited effects on these PRL variables. These findings show that MS does not lead to profound or persistent effects on behavioural flexibility but may produce more subtle sex-dependent effects on how feedback is processed.

Using a reversal learning task with both deterministic and probabilistic components, no difference was identified between the response to reversing contingencies in MS and control rats. While behavioural flexibility was hypothesised to be affected by early life stress, MS alone was not sufficient to produce lasting deficits in this measure of goal-directed behaviour. This conclusion is supported by our own recent work (Dutcher and colleagues (2023) and by Thomas et al. (2016) using a 4-choice odour discrimination task where deficits were reported in post-weaning juvenile rats but not adult rodents. Similarly, in humans, ELS was shown to impair cognitive flexibility in children (Harms et al. 2018) but not adults with a history of ELS (Demir-Lira et al. 2016; Franco and Knowlton 2023). Such findings indicate that the effects of ELS on behavioural flexibility are more apparent when testing occurs closer in temporal proximity of the early life stress (i.e. in children or younger animals) (Baudin et al. 2012; Harms et al. 2018). Thus, MS stress alone is not sufficient to persistently affect behavioural flexibility with compensation in the neural networks supporting flexible, goal-directed behaviour being apparently sufficient to offset any initial deficits.

Mimicking the endogenous stress response by administering corticosterone to adult rats was also insufficient to explicitly affect PRL based on conventional behavioural measures. Nevertheless, in control rats, corticosterone increased the modelled reward learning rate and side stickiness, decreased reinforcement sensitivity, while having no effect on reversal rate, win-stay or lose-shift responses. In contrast, administering corticosterone to MS rats decreased reward learning rate and side stickiness, increased reinforcement sensitivity and shortened the time for food collection. However, based on prior studies, the effects of exogenous stressors on reversal learning are broadly inconsistent, ranging from no effect (Wilkinson et al. 2020), to impaired reversal learning (Diederich and Trueblood 2018; Odland et al. 2023; Wilson et al. 2024), and increased response perseveration (Barfield and Gourley 2017). Nonetheless, our findings are reminiscent of Dutcher and colleagues (2023), who found that additional adult stress exacerbated already-longer response latencies on the PRL and impaired overall task performance for MS but not control subjects. Thus, building on the multi-hit hypothesis of stress sensitivity (Daskalakis et al. 2013), ELS changes the landscape for responses to future stressors. In contexts of acute stress, individuals become hypervigilant and expect a changing environment (Sara and Bouret 2012; Ross and Van Bockstaele 2021), the impaired behavioural flexibility seen in the present work may therefore represent an adaptive coping mechanism to blunt the high alert state when stressors are repeated. However, this may promote psychopathologies associated with impaired behavioural flexibility(Remijnse et al. 2006; Wilson et al. 2018; Wilkinson et al. 2021; Franco and Knowlton 2023).

Following the delivery of reward, MS increased the probability that the subsequent response would be directed to the same stimulus as the previous trial. This effect was most clearly expressed in males. The additional three-way interaction between sex, session and stress reflected a reduction in win-stay responding across sessions in female control animals, suggesting that MS may also affect the normal development or refinement of feedback-guided responding across repeated PRL sessions. These observations indicate that MS renders animals more sensitive to rewarding feedback in adulthood. We did not observe a sex dependent decrease in lose shift probability, which is in contrast to Dutcher and colleagues (2023), where the behavioural change was apparently driven by increased stickiness (Zühlsdorff et al. 2023). Of note, studies employing more severe stress procedures in mice report fewer win-stay responses following true rewards, and fewer shift responses following misleading losses, reflecting diminished feedback sensitivity, not stickiness (Bergamini et al. 2016). Studies with patients suffering from depression indicate an increased sensitivity to punishment on the PRL (Murphy et al. 2003; Taylor Tavares et al. 2008), as well as diminished estimates of value sensitivity (Mukherjee et al. 2020; Ogishima et al. 2020) and decreased win-stay responses following true wins (Dickstein et al. 2010). Wilkinson and colleagues (2021) also reported that both ELS in humans and acute recent stress resulted in decreased win-stay responding on the PRL, whereas lifetime stress in participants without ELS increased win-stay responding. Collectively, these findings in rodents and humans indicate that the severity and timing of stress in later life are key factors that shape the precise impacts of ELS on behavioural flexibility. More generally, the present study suggests that MS stress procedures have utility as an experimental approach to investigate latent vulnerability mechanisms that shape stress outcomes in later life.

In addition to conventional behavioural measures, we used computational modelling to extract several latent variables to inform reward and punishment learning rates, reinforcement sensitivity and stickiness. Behaviourally, MS males showed increased win-stay responding, suggesting greater persistence following rewarded feedback. In contrast, the reinforcement-learning model indicated increased punishment learning rate in MS males, with no corresponding change in reward learning rate, reinforcement sensitivity or side stickiness. This dissociation likely reflects the fact that summary measures such as win-stay and lose-shift capture overt trial-to-trial response patterns, whereas model parameters estimate latent updating processes across the task. Thus, the observed effect on win-stay behaviour should not be interpreted solely as increased reward learning. Rather, MS may alter the balance between overt response persistence and latent updating from negative outcomes. As the RL model was supported at the 75% HDI level, this interpretation should be viewed cautiously. We also investigated he involvement of NA and 5-HT systems in mediating the behavioural sequelae of MS. The β-2blocker propranolol produced a pattern consistent with increased response repetition or reduced choice variability. This compound increased reversal rate and win-stay responding while reducing lose-shift responding. At the highest dose tested it also increased perseverative responses. Thus, β-adrenergic blockade appeared to promote persistence with the currently favoured response. Such persistence may increase reversals once the correct option had been identified but impair rapid disengagement immediately after contingency reversal. Thus, propranolol slowed learning of new rules but reinforced responding according to the established rule; a result that replicates findings in humans where administration of propranolol reduced learning rate on a probabilistic associative learning task (Lawson et al. 2021). Indeed, other studies utilising clonidine, an α2-agonist, report similar effects on perseverative responding (Jahn et al. 2020) and behavioural flexibility (Ogg et al. 2023). In contrast, idazoxan, and α2-receptor antagonist, improved flexible responding (Ogg et al. 2023) while reboxetine, a selective NA reuptake inhibitor, decreased reversal rate and decreased win-stay responding (Wilkinson et al. 2020). Thus compounds which increase NA neurotransmission, either directly or indirectly, exert opposite effects to propranolol.

Prior studies have posited that LC NA signalling reflects volatility in the environment, including novelty, and the imperative to rapidly to update learnt associations (Bouret and Sara 2005; Dayan and Yu 2006; Avery et al. 2012). NA signalling also results in an increased readiness to abandon current learnt associations (Dayan and Yu 2006), decreased activation of fear memories (Santos-Mayo et al. 2022) and changes the balance of expectation and variability (Lawson et al. 2021). In patients with depression, the LC shows greater activation to both negatively valanced stimuli (Brendler et al. 2024). Using pupil size as a proxy for LC function, in the presence of greater stimulus noise and uncertainty, the LC encodes prediction errors and volatility in a PRL task; an effect which is abolished under propranolol (Lawson et al. 2021). Thus, blocking β-adrenergic signalling may reduce the tendency to update or explore alternative response-outcome contingencies, causing decisions to rely on previously learnt associations (Avery et al. 2012). However, despite evidence that MS affects the physiology and dendritic morphology of the LC (Swinny et al. 2010), this form of early life stress had surprisingly limited impact on the effects of propranolol on reversal learning.

Modulating 5-HT signalling with the SSRI citalopram also had minimal effects on PRL aside from evidence for an increase in punishment learning rate and reduced reinforcement sensitivity. Earlier studies with citalopram on this task reported either increased win-stay responding and a reduction in the number of trials to first reversal (Wilkinson et al. 2020) or an increased reversal rate and decreased lose-shift responding (Bari et al. 2010). In contrast, the serotonergic compounds escitalopram, venlafaxine and sertraline had no significant effect on PRL (Drozd et al. 2019; Wilkinson et al. 2020). In humans, acute citalopram administration increased lose-shift responding specifically following misleading punishment (Chamberlain et al. 2006; Skandali et al. 2018). However, artificially increasing 5-HT at the synapse using dietary tryptophan had no significant effect on PRL and DRL (Thirkettle et al. 2019; Kanen et al. 2021). Therefore, it appears that neither MS stress alone nor acute SSRI administration has robust effects on feedback sensitivity and reward learning with respect to reversal learning.

Several limitations should be considered. First, the effects of MS on PRL were selective and sex-dependent, and most conventional task-performance measures were unaffected. The findings should therefore not be interpreted as evidence for a broad impairment in behavioural flexibility. Second, the pharmacological manipulations were systemic and acute, so the observed effects cannot be attributed specifically to LC–PFC noradrenergic or raphe serotonergic circuits nor capture the effects of chronic modulation relevant to clinical treatment. Third, several computational effects were supported at a 75% HDI threshold and should be considered suggestive rather than definitive; this may be compounded by the relatively limited number of model checks, particularly the lack of parameter identifiability checks, performed beyond model comparisons. Finally, although corticosterone provides a useful neuroendocrine manipulation, it does not reproduce the full physiological, behavioural and contextual repertoire of psychological stress.

In conclusion, the present study highlights the experimental utility of MS to investigate ELS, specifically latent vulnerability mechanisms that affect feedback sensitivity on a PRL task. Our findings indicate that MS has persistent but sexually dimorphic effects on the sensitivity of reward and feedback processing. However, despite evidence of persistent interactive effects of MS on the serotonergic and noradrenergic systems, MS did not affect the modulation of serial probabilistic learning by propranolol and citalopram. Such null findings potentially point to the importance of multiple stress hits across the lifespan in the manifestation of psychopathologies involving impaired behavioural flexibility.

## Acknowledgements

We are grateful to Rudolf Cardinal for his contribution to the computational modelling component of this research and Amy Milton as the Home Office project licence holder.

## Author contributions

OS and JWD together designed the study; OS, LJFW and JC conducted the measurements; OS and KZ performed data analyses; OS and LJFW wrote the first draft of the manuscript, which was edited by JWD, CVS, MCDB and RPL; OS, LJFWV and JWD drafted the revised manuscript; all authors approved the initially submitted and revised manuscripts.

## Data availability

Data are available upon reasonable request from the corresponding author.

## Ethical approval

Experiments were conducted in accordance with the Animals (Scientific procedures) Act, 1983, Amendment Regulations (2012) on project licence PA9FBFA9F following ethical review by the University of Cambridge Animal Welfare and Ethical Review Body (AWERB).

## Open Access

This article is licensed under a Creative Commons Attribution 4.0 International License, which permits use, sharing, adaptation, distribution and reproduction in any medium or format, as long as you give appropriate credit to the original author(s) and the source, provide a link to the Creative Commons licence, and indicate if changes were made. The images or other third-party material in this article are included in the article’s Creative Commons licence, unless indicated otherwise in a credit line to the material. If material is not included in the article’s Creative Commons licence and your intended use is not permitted by statutory regulation or exceeds the permitted use, you will need to obtain permission directly from the copyright holder. To view a copy of this licence, visit http://creativecommons.org/licenses/by/4.0/.

## References

Agnew-Blais J, Danese A (2016) Childhood maltreatment and unfavourable clinical outcomes in bipolar disorder: a systematic review and meta-analysis. Lancet Psychiatry 3:342–349. 10.1016/S2215-0366(15)00544-1

Avery MC, Nitz DA, Chiba AA, Krichmar JL (2012) Simulation of cholinergic and noradrenergic modulation of behavior in uncertain environments. Front Comput Neurosci 6:. 10.3389/fncom.2012.00005

Barfield ET, Gourley SL (2017) Adolescent Corticosterone and TrkB Pharmaco-Manipulations Sex-Dependently Impact Instrumental Reversal Learning Later in Life. Front Behav Neurosci Volume 11–2017: 10.3389/fnbeh.2017.00237

Bari A, Theobald DE, Caprioli D, et al (2010) Serotonin modulates sensitivity to reward and negative feedback in a probabilistic reversal learning task in rats. Neuropsychopharmacology 35:1290–1301. 10.1038/npp.2009.233

Baudin A, Blot K, Verney C, et al (2012) Maternal deprivation induces deficits in temporal memory and cognitive flexibility and exaggerates synaptic plasticity in the rat medial prefrontal cortex. Neurobiol Learn Mem 98:207–214. 10.1016/j.nlm.2012.08.004

Bergamini G, Cathomas F, Auer S, et al (2016) Mouse psychosocial stress reduces motivation and cognitive function in operant reward tests: A model for reward pathology with effects of agomelatine. European Neuropsychopharmacology 26:1448–1464. 10.1016/j.euroneuro.2016.06.009

Bouret S, Sara SJ (2005) Network reset: a simplified overarching theory of locus coeruleus noradrenaline function. Trends Neurosci 28:574–582. 10.1016/j.tins.2005.09.002

Brendler A, Schneider M, Elbau IG, et al (2024) Assessing hypo-arousal during reward anticipation with pupillometry in patients with major depressive disorder: replication and correlations with anhedonia. Sci Rep 14:344. 10.1038/s41598-023-48792-0

Browning M, Behrens TE, Jocham G, et al (2015) Anxious individuals have difficulty learning the causal statistics of aversive environments. Nat Neurosci 18:590–596. 10.1038/nn.3961

Bryce CA, Floresco SB (2021) Central CRF and acute stress differentially modulate probabilistic reversal learning in male and female rats. Behavioural Brain Research 397:112929. 10.1016/j.bbr.2020.112929

Carpenter B, Gelman A, Hoffman MD, et al (2017) Stan: A Probabilistic Programming Language. J Stat Softw 76:1–32. 10.18637/jss.v076.i01

Cella M, Dymond S, Cooper A (2010) Impaired flexible decision-making in major depressive disorder. J Affect Disord 124:207–210. 10.1016/j.jad.2009.11.013

Chamberlain SR, Müller U, Blackwell AD, et al (2006) Neurochemical modulation of response inhibition and probabilistic learning in humans. Science 311:861–863. 10.1126/science.1121218

Cohen RA, Hitsman BL, Paul RH, et al (2006) Early Life Stress and Adult Emotional Experience: An International Perspective. The International Journal of Psychiatry in Medicine 36:35–52. 10.2190/5R62-9PQY-0NEL-TLPA

Daskalakis NP, Bagot RC, Parker KJ, et al (2013) The three-hit concept of vulnerability and resilience: Toward understanding adaptation to early-life adversity outcome. Psychoneuroendocrinology 38:1858–1873. 10.1016/j.psyneuen.2013.06.008

Dayan P, Yu AJ (2006) Phasic norepinephrine: A neural interrupt signal for unexpected events. Network: Computation in Neural Systems 17:335–350. 10.1080/09548980601004024

Delcourte S, Etievant A, Haddjeri N (2021) Chapter 2 - Role of central serotonin and noradrenaline interactions in the antidepressants’ action: Electrophysiological and neurochemical evidence. In: Di Giovanni G, De Deurwaerdere P (eds) Progress in Brain Research. Elsevier, pp 7–81

Demir-Lira ÖE, Voss JL, O’Neil JT, et al (2016) Early-life stress exposure associated with altered prefrontal resting-state fMRI connectivity in young children. Dev Cogn Neurosci 19:107–114. 10.1016/j.dcn.2016.02.003

Dickstein DP, Finger EC, Brotman MA, et al (2010) Impaired probabilistic reversal learning in youths with mood and anxiety disorders. Psychol Med 40:1089–1100. 10.1017/S0033291709991462

Diederich A, Trueblood JS (2018) A dynamic dual process model of risky decision making. Psychol Rev 125:270–292. 10.1037/rev0000087

Drozd R, Rychlik M, Fijalkowska A, Rygula R (2019) Effects of cognitive judgement bias and acute antidepressant treatment on sensitivity to feedback and cognitive flexibility in the rat version of the probabilistic reversal-learning test. Behavioural Brain Research 359:619–629. 10.1016/j.bbr.2018.10.003

Dutcher EG, Lopez-Cruz L, Pama EAC, et al (2023) Early-life stress biases responding to negative feedback and increases amygdala volume and vulnerability to later-life stress. Transl Psychiatry 13:81. 10.1038/s41398-023-02385-7

Elliott R, Rubinsztein JS, Sahakian BJ, Dolan RJ (2002) The Neural Basis of Mood-Congruent Processing Biases in Depression. Arch Gen Psychiatry 59:597. 10.1001/archpsyc.59.7.597

Elliott R, Sahakian BJ, Herrod JJ, et al (1997) Abnormal response to negative feedback in unipolar depression: evidence for a diagnosis specific impairment. Journal of Neurology, Neurosurgery & Psychiatry 63:74. 10.1136/jnnp.63.1.74

Fóscolo DRC, Lima PMA, Rodovalho G V, Coimbra CC (2022) Early maternal separation alters the activation of stress-responsive brain areas in adulthood. Neurosci Lett 771:136464. 10.1016/j.neulet.2022.136464

Franco CY, Knowlton BJ (2023) Effects of early-life stress on probabilistic reversal learning and response perseverance in young adults. Neurobiol Learn Mem 205:107839. 10.1016/j.nlm.2023.107839

Grella SL, Gomes SM, Lackie RE, et al (2021) Norepinephrine as a spatial memory reset signal. Behavioural Pharmacology 32:

Gronau QF, Sarafoglou A, Matzke D, et al (2017) A tutorial on bridge sampling. J Math Psychol 81:80–97. 10.1016/j.jmp.2017.09.005

Harms MB, Shannon Bowen KE, Hanson JL, Pollak SD (2018) Instrumental learning and cognitive flexibility processes are impaired in children exposed to early life stress. Dev Sci 21:e12596. 10.1111/desc.12596

Hauser TU, Allen M, Purg N, et al (2017) Noradrenaline blockade specifically enhances metacognitive performance. Elife 6:e24901. 10.7554/eLife.24901

Hecht PM, Will MJ, Schachtman TR, et al (2014) Beta-adrenergic antagonist effects on a novel cognitive flexibility task in rodents. Behavioural Brain Research 260:148–154. 10.1016/j.bbr.2013.11.041

Hernández-Pérez OR, Hernández VS, Nava-Kopp AT, et al (2019) A Synaptically Connected Hypothalamic Magnocellular Vasopressin-Locus Coeruleus Neuronal Circuit and Its Plasticity in Response to Emotional and Physiological Stress. Front Neurosci Volume 13–2019:

Itoi K, Sugimoto N (2010) The Brainstem Noradrenergic Systems in Stress, Anxiety and Depression. J Neuroendocrinol 22:355–361. 10.1111/j.1365-2826.2010.01988.x

Jahn CI, Varazzani C, Sallet J, et al (2020) Noradrenergic But Not Dopaminergic Neurons Signal Task State Changes and Predict Reengagement After a Failure. Cerebral Cortex 30:4979–4994. 10.1093/cercor/bhaa089

Kanen JW, Apergis-Schoute AM, Yellowlees R, et al (2021) Serotonin depletion impairs both Pavlovian and instrumental reversal learning in healthy humans. Mol Psychiatry 26:7200–7210. 10.1038/s41380-021-01240-9

Lawson RP, Bisby J, Nord CL, et al (2021) The Computational, Pharmacological, and Physiological Determinants of Sensory Learning under Uncertainty. Current Biology 31:163–172.e4. 10.1016/j.cub.2020.10.043

Lenth R V (2024) emmeans: Estimated marginal means, aka least-squares means

Lissek S (2012) TOWARD AN ACCOUNT OF CLINICAL ANXIETY PREDICATED ON BASIC, NEURALLY MAPPED MECHANISMS OF PAVLOVIAN FEAR-LEARNING: THE CASE FOR CONDITIONED OVERGENERALIZATION. Depress Anxiety 29:257–263. 10.1002/da.21922

Llorente R, O’Shea E, Gutierrez-Lopez MD, et al (2010) Sex-dependent maternal deprivation effects on brain monoamine content in adolescent rats. Neurosci Lett 479:112–117. 10.1016/j.neulet.2010.05.039

Matthews K, Robbins TW (2003) Early experience as a determinant of adult behavioural responses to reward: the effects of repeated maternal separation in the rat. Neurosci Biobehav Rev 27:45–55. 10.1016/S0149-7634(03)00008-3

McCall JG, Siuda ER, Bhatti DL, et al (2017) Locus coeruleus to basolateral amygdala noradrenergic projections promote anxiety-like behavior. Elife 6:e18247. 10.7554/eLife.18247

Morris LS, McCall JG, Charney DS, Murrough JW (2020) The role of the locus coeruleus in the generation of pathological anxiety. Brain Neurosci Adv 4:239821282093032. 10.1177/2398212820930321

Mukherjee D, Filipowicz ALS, Vo K, et al (2020) Reward and punishment reversal-learning in major depressive disorder. J Abnorm Psychol 129:810–823. 10.1037/abn0000641

Murphy FC, Michael A, Robbins TW, Sahakian BJ (2003) Neuropsychological impairment in patients with major depressive disorder: the effects of feedback on task performance. Psychol Med 33:455–467. DOI: 10.1017/S0033291702007018

Nanni V, Uher R, Danese A (2012) Childhood Maltreatment Predicts Unfavorable Course of Illness and Treatment Outcome in Depression: A Meta-Analysis. American Journal of Psychiatry 169:141–151. 10.1176/appi.ajp.2011.11020335

Nishi M, Horii-Hayashi N, Sasagawa T (2014) Effects of early life adverse experiences on the brain: implications from maternal separation models in rodents. Front Neurosci Volume 8–2014:

Odland AU, Sandahl R, Andreasen JT (2023) Chronic corticosterone improves perseverative behavior in mice during sequential reversal learning. Behavioural Brain Research 450:114479. 10.1016/j.bbr.2023.114479

Ogg MC, Franks HT, Lansdell BJ, et al (2023) Locus Coeruleus Norepinephrine Neurons Facilitate Orbitofrontal Cortex Remapping and Behavioral Flexibility

Ogishima H, Maeda S, Tanaka Y, Shimada H (2020) Effects of Depressive Symptoms, Feelings, and Interoception on Reward-Based Decision-Making: Investigation Using Reinforcement Learning Model. Brain Sci 10:. 10.3390/brainsci10080508

Ohta K, Miki T, Warita K, et al (2014) Prolonged maternal separation disturbs the serotonergic system during early brain development. International Journal of Developmental Neuroscience 33:15–21. 10.1016/j.ijdevneu.2013.10.007

Remijnse PL, Nielen MMA, Van Balkom AJLM, et al (2006) Reduced Orbitofrontal-Striatal Activity on a Reversal Learning Task in Obsessive-Compulsive Disorder. Arch Gen Psychiatry 63:1225. 10.1001/archpsyc.63.11.1225

Rentesi G, Antoniou K, Marselos M, et al (2010) Long-term consequences of early maternal deprivation in serotonergic activity and HPA function in adult rat. Neurosci Lett 480:7–11. 10.1016/j.neulet.2010.04.054

Rex A, Voigt JP, Voits M, Fink H (1998) Pharmacological Evaluation of a Modified Open-Field Test Sensitive to Anxiolytic Drugs. Pharmacol Biochem Behav 59:677–683. 10.1016/S0091-3057(97)00461-9

Richelson E (1994) Pharmacology of Antidepressants-Characteristics of the Ideal Drug. Mayo Clin Proc 69:1069–1081. 10.1016/S0025-6196(12)61375-5

Rorabaugh JM, Chalermpalanupap T, Botz-Zapp CA, et al (2017) Chemogenetic locus coeruleus activation restores reversal learning in a rat model of Alzheimer’s disease. Brain 140:3023–3038. 10.1093/brain/awx232

Ross JA, Van Bockstaele EJ (2021) The Locus Coeruleus-Norepinephrine System in Stress and Arousal: Unraveling Historical, Current, and Future Perspectives. Front Psychiatry Volume 11–2020:

Santos-Mayo A, De Echegaray J, Moratti S (2022) Conditioned up and down modulations of short latency gamma band oscillations in visual cortex during fear learning in humans. Sci Rep 12:2652. 10.1038/s41598-022-06596-8

Sara SJ, Bouret S (2012) Orienting and Reorienting: The Locus Coeruleus Mediates Cognition through Arousal. Neuron 76:130–141. 10.1016/j.neuron.2012.09.011

Sara SJ, Vankov A, Hervé A (1994) Locus coeruleus-evoked responses in behaving rats: A clue to the role of noradrenaline in memory. Brain Res Bull 35:457–465. 10.1016/0361-9230(94)90159-7

Schick A, Adam R, Vollmayr B, et al (2015) Neural correlates of valence generalization in an affective conditioning paradigm. Behavioural Brain Research 292:147–156. 10.1016/j.bbr.2015.06.009

Skandali N, Rowe JB, Voon V, et al (2018) Dissociable effects of acute SSRI (escitalopram) on executive, learning and emotional functions in healthy humans. Neuropsychopharmacology 43:2645–2651. 10.1038/s41386-018-0229-z

Smith SM, Vale WW (2006) The role of the hypothalamic-pituitary-adrenal axis in neuroendocrine responses to stress. Dialogues Clin Neurosci 8:383–395. 10.31887/DCNS.2006.8.4/ssmith

Soubrié P (1986) Neurones sérotoninergiques et comportement [Serotonergic neurons and behavior]. J Pharmacol 17:107–12

Swinny JD, O’Farrell E, Bingham BC, et al (2010) Neonatal rearing conditions distinctly shape locus coeruleus neuronal activity, dendritic arborization, and sensitivity to corticotrophin-releasing factor. Int J Neuropsychopharmacol 13:515–525. 10.1017/S146114570999037X

Taylor Tavares J V, Clark L, Furey ML, et al (2008) Neural basis of abnormal response to negative feedback in unmedicated mood disorders. Neuroimage 42:1118–1126. 10.1016/j.neuroimage.2008.05.049

Thirkettle M, Barker L-M, Gallagher T, et al (2019) Dissociable Effects of Tryptophan Supplementation on Negative Feedback Sensitivity and Reversal Learning. Front Behav Neurosci 13:127. 10.3389/fnbeh.2019.00127

Thomas AW, Caporale N, Wu C, Wilbrecht L (2016) Early maternal separation impacts cognitive flexibility at the age of first independence in mice. Dev Cogn Neurosci 18:49–56. 10.1016/j.dcn.2015.09.005

Vahid-Ansari F, Albert PR (2021) Rewiring of the Serotonin System in Major Depression. Front Psychiatry Volume 12-2021:

Wang L, Dai Z, Peng H, et al (2014) Overlapping and segregated resting-state functional connectivity in patients with major depressive disorder with and without childhood neglect. Hum Brain Mapp 35:1154–1166. 10.1002/hbm.22241

Wilkinson MP, Grogan JP, Mellor JR, Robinson ESJ (2020) Comparison of conventional and rapid-acting antidepressants in a rodent probabilistic reversal learning task. Brain Neurosci Adv 4:239821282090717. 10.1177/2398212820907177

Wilkinson MP, Slaney CL, Mellor JR, Robinson ESJ (2021) Investigation of reward learning and feedback sensitivity in non-clinical participants with a history of early life stress. PLoS One 16:e0260444. 10.1371/journal.pone.0260444

Wilson C, Hannan AJ, Renoir T (2024) Serotonergic agonism and pharmacologically-induced adolescent stress cause operant-based learning deficits in mice. Neuropharmacology 244:109801. 10.1016/j.neuropharm.2023.109801

Wilson CG, Nusbaum AT, Whitney P, Hinson JM (2018) Trait anxiety impairs cognitive flexibility when overcoming a task acquired response and a preexisting bias. PLoS One 13:e0204694. 10.1371/journal.pone.0204694

Wong DT, Threlkeld PG, Robertson DW (1991) Affinities of fluoxetine, its enantiomers, and other inhibitors of serotonin uptake for subtypes of serotonin receptors. Neuropsychopharmacology 5:43–47

Yu AJ, Dayan P (2005) Uncertainty, neuromodulation, and attention. Neuron 46:681–692. 10.1016/j.neuron.2005.04.026

Zaidi S, Atrooz F, Valdez D, et al (2020) Protective effect of propranolol and nadolol on social defeat-induced behavioral impairments in rats. Neurosci Lett 725:134892. 10.1016/j.neulet.2020.134892

Zühlsdorff K, López-Cruz L, Dutcher EG, et al (2023) Sex-dependent effects of early life stress on reinforcement learning and limbic cortico-striatal functional connectivity. Neurobiol Stress 22:100507. 10.1016/j.ynstr.2022.100507

